# Condensation of the fusion focus by the intrinsically disordered region of the formin Fus1 is essential for cell-cell fusion

**DOI:** 10.1101/2022.05.05.490810

**Authors:** Ingrid Billault-Chaumartin, Olivia Muriel, Laetitia Michon, Sophie G Martin

## Abstract

Spatial accumulation of secretory vesicles underlies various cellular processes, such as neurotransmitter release at neuronal synapses [1], hyphal steering in filamentous fungi [2, 3], and local cell wall digestion preceding the fusion of yeast gametes [4]. Secretory vesicles transported on actin filaments by myosin V motors form clusters that serve as pool for local content release. During fission yeast *Schizosaccharomyces pombe* gamete fusion, the actin fusion focus assembled by the formin Fus1 concentrates secretory vesicles carrying cell wall digestive enzymes [5-7]. Focus position and coalescence are controlled by local signalling and actin-binding proteins to prevent inappropriate cell wall digestion that would cause lysis [6, 8-10], but the mechanisms of focusing have been elusive. Here, we show that the regulatory N-terminus of Fus1 contains an intrinsically disordered region (IDR) that mediates Fus1 condensation in vivo and forms dense assemblies that exclude other macromolecules. Fus1 lacking its IDR fails to condense in a tight focus and causes cell lysis during attempted cell fusion. Remarkably, replacement of Fus1 IDR with a heterologous low-complexity region that forms liquid condensates fully restores Fus1 condensation and function. By contrast, replacement of Fus1 IDR with a domain that forms more stable oligomers restores condensation but poorly supports cell fusion, suggesting that condensation is tuned to yield a structure selectively permeable for secretory vesicles. We propose that condensation of actin structures by an intrinsically disordered region may be a general mechanism for actin network organisation and the selective local concentration of secretory vesicles.

## Results and discussion

### Fus1N has localization and self-association properties

We previously showed that Fus1 function during cell fusion cannot be fulfilled by either of the other two fission yeast formins, For3 and Cdc12 [11]. However, chimeric proteins containing the Fus1 regulatory N-terminus (called Fus1N below) and the actin nucleating FH1-FH2 domains of For3 or Cdc12 partially rescued *fus1Δ* defects, indicating an important role for Fus1N. To test the roles of Fus1N, we replaced it with the N-termini of For3 and Cdc12 (Figure 1A). Neither Cdc12N-Fus1C nor For3N-Fus1C supported cell fusion, while the control Fus1N-Fus1C was fully functional (Figure 1B-C). While Cdc12N-Fus1C failed to localize, For3N-Fus1C localized to the region of contact between the two cells, albeit over a wider zone than Fus1 (Figure 1B). These results suggest that localization to the shmoo tip is not the only Fus1N property required for cell fusion.

**Figure 1.**
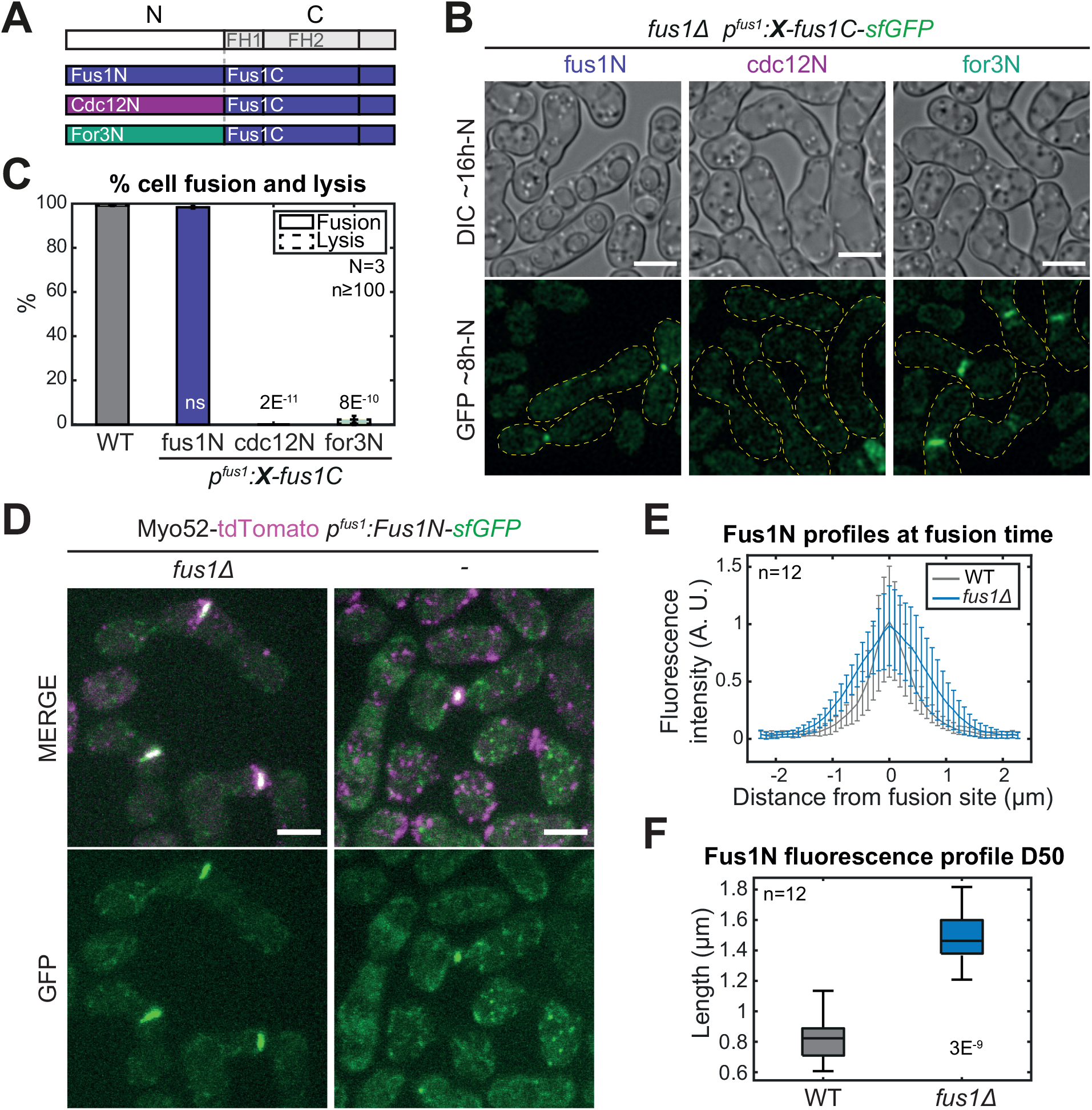
Fus1N is essential for fusion and associates with the fusion focus. **A**. Formin chimeras tagged C-terminally with sfGFP. **B**. DIC and GFP images ∼16h and ∼8h post starvation of *fus1Δ* cells expressing the chimeric formins shown in (A). Yellow dashed lines outline mating pairs. **C**. Percentage of cell pair fusion and lysis 24h post starvation in WT and strains as in (B). **D**. Merge and GFP images ∼8h post starvation of Myo52-tdTomato and Fus1N-sfGFP (Fus1^1-792^) expressed in *fus1Δ* or WT cells. **E**. Normalized Fus1N-sfGFP fluorescence profiles perpendicular to the mating pair axis at the time of cell fusion, in strains as in (D). **F**. Width at half maximum of the profiles shown in (E). Bars are 5µm. p-values relative to WT.

A Fus1N fragment localized to the region of cell-cell contact, as previously reported [12], in both *fus1Δ* and wildtype cells. However, its precise distribution was different in the two backgrounds. In *fus1Δ*, Fus1N decorated the entire cell-cell contact area, like Myo52 (Figure 1D). When expressed in addition to endogenous WT Fus1, Fus1N localized in a restricted focus coincident with the Myo52 marker (Figure 1D). Measurements of Fus1N distribution along the plasma membrane showed a nearly two-fold narrower distribution in presence of WT Fus1 (Figure 1E-F). These results indicate that Fus1N associates with the fusion focus, perhaps through association with Fus1 itself.

### Fus1N contains separable tip-localisation and cluster-forming properties

Fus1N (aa 1-792) contains a GBD/FH3 domain that mediates localization [12], followed by an intrinsically disordered region (IDR), as predicted by tools such as ODiNPred [13], IUPred3 [14] and PONDR [15] (Figure 2A). Fus1N expressed in interphase cells exhibited a dual localization to cell tips and cytosolic clusters (Figure 2A-B). Shortening Fus1N from the N-terminus led to progressive loss of cell tip localisation (Fus1N^93-792^, Fus1N^140-792^ and Fus1N^191-792^; Figure 2A-B). Truncation of Fus1N IDR from its C-terminus led to a progressive loss of cytosolic clusters (Fus1N^1-730^ and Fus1N^1-500^; Figure 2A-B). When shortened from both ends, Fus1N lost both localizations (Fus1N^431-755^; Figure 2A-B). Thus, Fus1 N-terminal extremity and IDR provide distinct localization and clustering functions.

**Figure 2.**
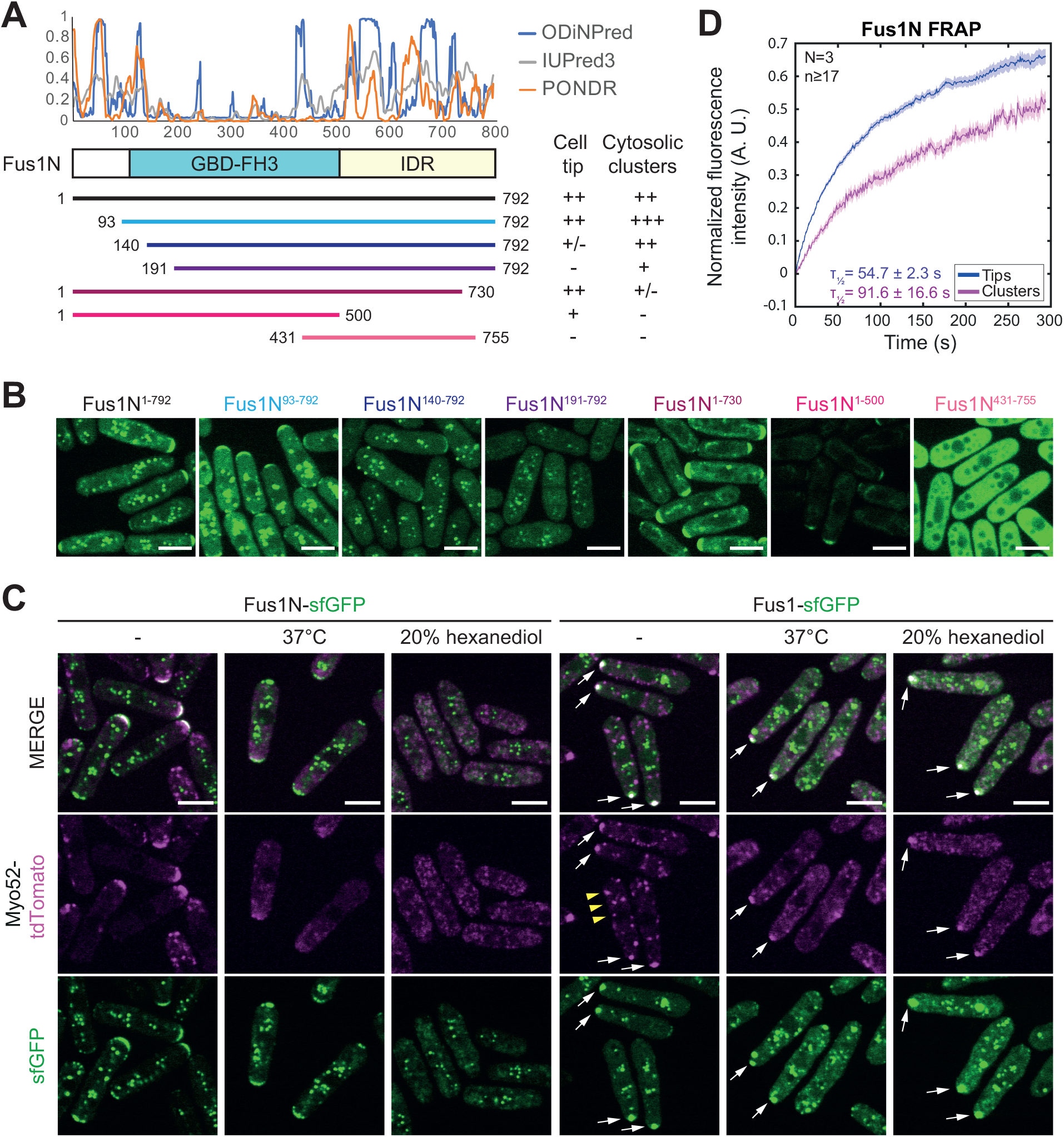
Fus1N contains separable tip-localization and cluster-forming determinants. **A**. Scheme of Fus1N with predicted domain organization. The top graph shows the disorder index of 3 prediction tools [13-15]. Fragments were C-terminally tagged with sfGFP. Localization summary is shown on the right. **B**. GFP fluorescence images of constructs as in (A). **C**. Fluorescence images of interphase cells expressing Myo52-tdTomato and either (left) Fus1N-sfGFP (Fus1^1-792^) or (right) full length Fus1-sfGFP from the *nmt1* promotor. Cells were either untreated (left), grown for 6h at 37°C and imaged at 40°C (middle), or treated with 20% 1,6-hexanediol for 5 minutes (right). White arrows mark resistant fusion focus-like structure; yellow arrowheads indicate labile Myo52 dots. **D**. Average Fus1N FRAP recovery curves normalized to pre-bleach values of cells as in (C). The mean recovery half-time and standard deviation from 3 independent experiments is indicated. N=3 experiments, with n>17 cells each (n>54 cells in total). Shaded area shows the standard error. Bars are 5µm.

In these experiments, we noticed that Fus1N expression modified cellular growth patterns. Whereas WT cells show bipolar growth and activate Cdc42 GTPase and actin assembly at both cell poles, Fus1N^1-792^ often localized to a single cell tip, where it colocalized with CRIB labelling Cdc42-GTP [16] and the myosin V Myo52 (Figure S1A, 2C). These markers were largely absent from the second cell tip. They also did not colocalize with Fus1N cytosolic clusters. By contrast, Tea1, a cell end marker transported by microtubules [17] was more prominently enriched at the other cell tip (Figure S1A), a localization previously described in monopolar *tea4Δ* mutants [18]. The monopolar phenotype was more prominent in cells expressing Fus1N^1-730^, which almost exclusively localizes to cell tips, but absent in cells expressing Fus1^93-792^, which partitions mainly to cytosolic clusters (Figures 2B and S1), suggesting that the phenotype results from Fus1N presence at the cell tip. While we do not fully understand how Fus1N induces monopolarity, one possibility is that this fragment concentrates growth components at a single cell tip, preventing initiation of cell growth at the second cell pole.

To better understand the nature of the Fus1N clusters, we subjected them, as well as Myo52 tagged in the same cell, to high temperature or 1,6-hexanediol treatments, known to compromise weak interactions [19]. Exposure of cells to 37°C severely disturbed Myo52 localization but left Fus1N localization unaffected (Figure 2C, left). Treatment of cells with 20% 1,6-hexanediol, an aliphatic alcohol that interferes with hydrophobic interactions and is widely used for disrupting liquid-liquid phase separated (LLPS) condensates [20], dissipated the cell tip localization of Fus1N and Myo52, and reduced Fus1N clusters, although they were still present, suggesting a solid core (Figure 2C, left). Fluorescence recovery after photobleaching (FRAP) experiments further suggested a higher stability of Fus1N in cytosolic clusters than at cell tips: in cytosolic clusters, only about 50% of the Fus1N signal was mobile and recovered more slowly than the larger mobile pool at cell poles (Figure 2D). Taken together, these experiments indicate that Fus1N forms resistant assemblies in cells.

Full-length Fus1 expressed in interphase cells formed both cytosolic clusters that did not colocalize with Myo52, like the Fus1N clusters, and a prominent focus that recruited Myo52, like the fusion focus during sexual reproduction (Figure 2C, right). This focus localized preferentially at one cell pole, which was thinner, or at the division site, occasionally leading to cell lysis after division (Movie S1). Thus, Fus1 is active when expressed in mitotic cells and likely concentrates secretion over a narrow region, leading to cell thinning and lysis. High temperature and 1,6-hexanediol treatments increased cytosolic signals and disassembled the small, likely vesicle-associated Myo52 dots (yellow arrowheads), but disrupted neither Fus1 cytosolic clusters nor the larger focus of Fus1 and Myo52 (white arrows). This suggests that recruitment of Myo52 upon actin polymerisation by Fus1 traps the motor protein (and likely associated vesicles) in the Fus1 structure.

### Fus1 foci are zones of ribosome exclusion

In correlative light-electron microscopy (CLEM) studies, we previously reported that fusion foci accumulate secretory vesicles but exclude ribosomes and other organelles [7], suggesting they represent membrane-less organelles. Ribosomes may be largely excluded due to space occupancy by secretory vesicles and associated factors. Alternatively, the fusion focus may exclude large macro-molecular components independently of secretory vesicles. To probe this question, we acquired CLEM-tomograms of Fus1-sfGFP labelled cell pairs lacking the capping protein β subunit Acp2, in which Fus1 accumulates excessively at the fusion focus but the recruitment of Myo52 and secretory vesicle markers is reduced by about two-fold [21]. Indeed, ultrastructure of the fusion site in *acp2Δ* showed reduced secretory vesicle recruitment relative to WT (Figure 3A-C). However, the zone of *acp2Δ* cell-cell contact showed a region devoid of ribosomes, at least as large as that observed in WT cells, indicating local macromolecular exclusion by molecular crowding and/or actin assembly independently of the presence of secretory vesicles (Figure 3A-B, 3D).

**Figure 3.**
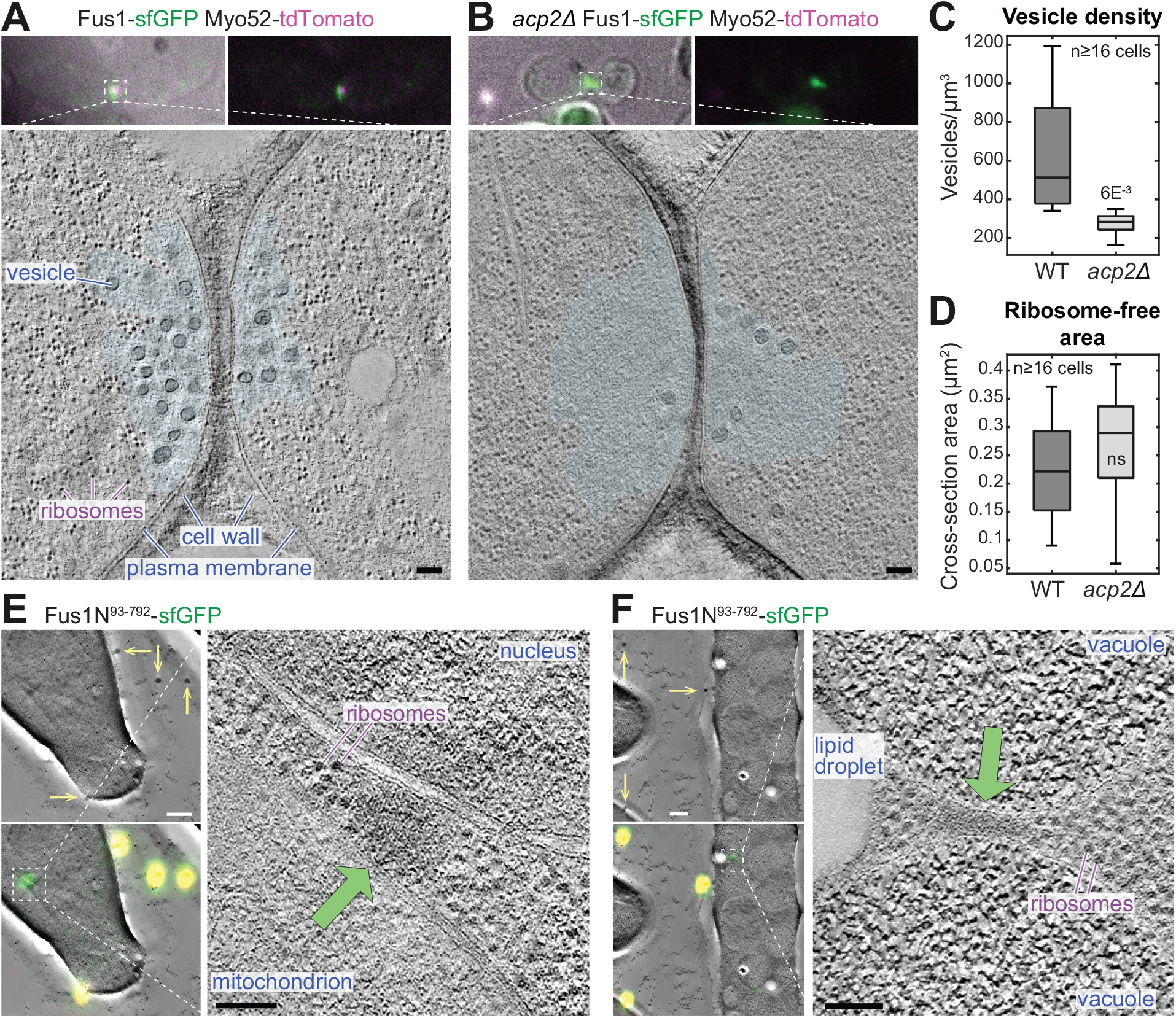
Fus1 assemblies exclude ribosomes. **A-B**. Virtual z-slices through electron tomograms taken at the contact site of WT and *acp2Δ* cell pairs during the fusion process. The transparent cyan shape outlines regions almost completely devoid of ribosomes. Images on top show transmitted light image and fluorescence of Fus1-sfGFP (green) and Myo52-tdTomato (magenta). **C**. Density of vesicles within a half-cylinder of diameter 1 µm centered at the cells’ contact zone. The p-value is shown. **D**. Cross-section area of the ribosome-free zone in cells as in (A-B). **E-F**. Virtual z-slices through electron tomograms at the position of Fus1N^93-792^-sfGFP signal in vegetative cells. Images on the left show low-magnification tomograms with and without the correlated fluorescence image of Fus1N^93-792^-sfGFP (green) and fiducial beads (yellow and arrows). Scale bars are 100 nm, except for (D-E, left: 500 nm).

To test whether Fus1 condensation can promote macromolecular exclusion independently of actin assembly, we further acquired CLEM-tomograms of Fus1N^93-792^, which lacks actin assembly capacity and forms prominent cytosolic clusters. In 26 of 30 tomograms, the Fus1N^93-792^-sfGFP fluorescence signal was positioned within 100nm (corresponding to the precision of the correlation) of a 100-300nm-wide cytosolic region devoid of ribosomes (Figure 3E-F). In 23 of these, the region was also darker than the surrounding cytosol. Thus, independently of actin assembly, Fus1N forms large, dense structures that exclude other macromolecules. We postulate that it similarly contributes to molecular crowding in the fusion focus.

### Fus1 IDR concentrates Fus1 and is essential for fusion

To test the functional relevance of Fus1 self-assembly, we deleted the IDR from full-length Fus1 expressed from the endogenous locus, constructing three deletions of increasing length. Each mutant protein localized correctly to the site of cell-cell contact but distributed over a wider area than WT Fus1 (Figure 4A-B, first four columns). This increased width was most extensive for the mutant with the longest deletion, Fus1^Δ492-791^ (Figure 4B). Thus, Fus1 IDR strongly contributes to the concentration of Fus1 in a focus.

**Figure 4.**
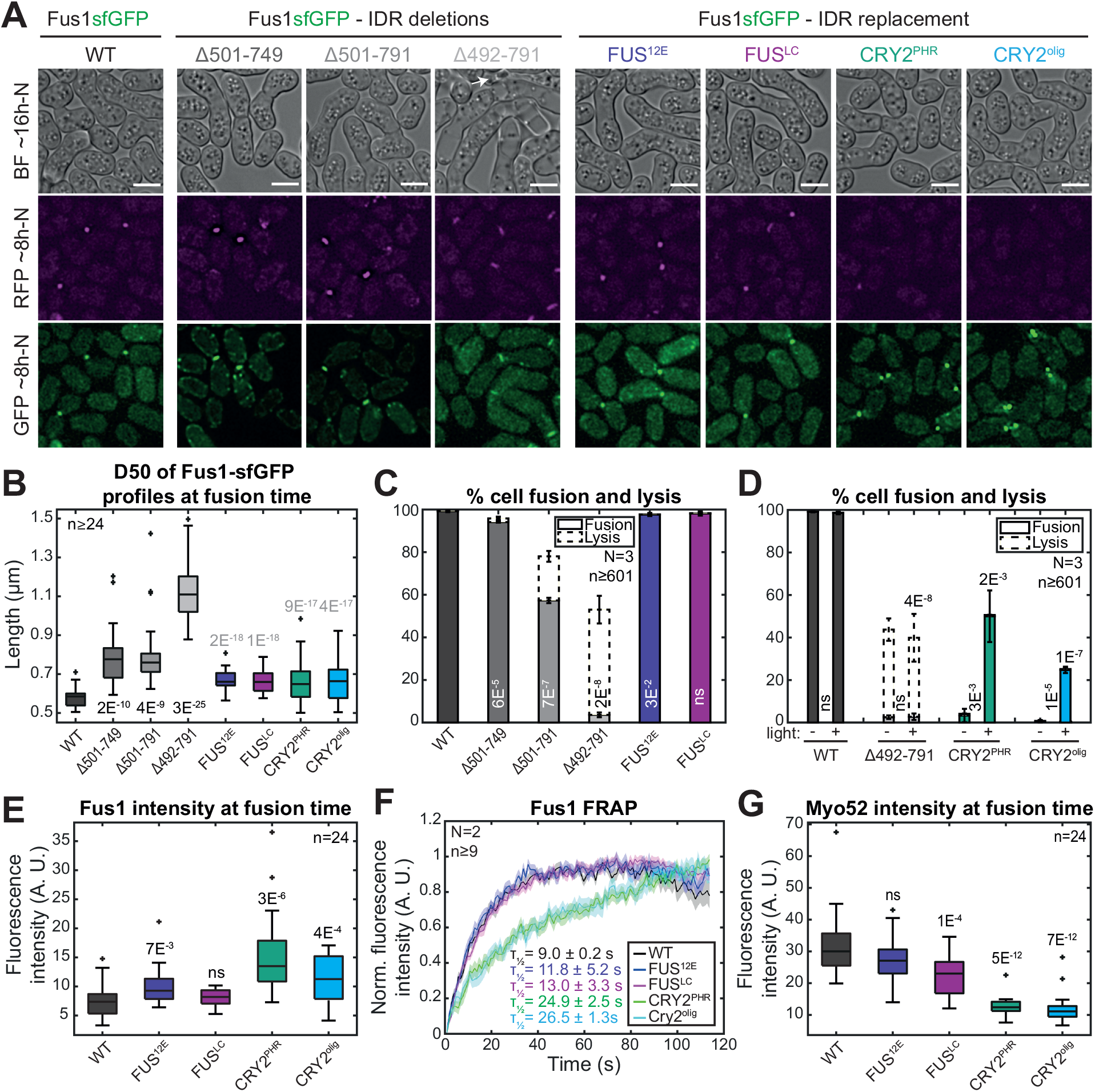
Fus1 IDR is essential for fusion and functions as a clustering device. **A**. DIC and fluorescence images ∼16h and ∼8h post starvation of cells expressing Myo52-tdTomato and the indicated Fus1-sfGFP allele from the native *fus1* locus. FUS and CRY2 variants were introduced in Fus1^Δ492-791^. Cells were exposed to blue light every 5 minutes for several hours. The white arrow points to a lysed pair. Lysis is under-represented, as it mostly happens at later timepoints. **B**. Width at half maximum of GFP-fluorescence profiles in strains as in (A), similarly to Figure 1E-F. **C**. Percentage of cell pair fusion and lysis 24h post starvation in strains with Fus1 or Fus1-FUS alleles as in (A). **D**. Percentage of cell pair fusion and lysis 24h post starvation under continuous white light exposure (+) or in the dark (-) in strains with Fus1 or Fus1-CRY2 alleles as in (A). **E**. Boxplot of Fus1 fluorescence intensity at the fusion focus at fusion time. **F**. Average Fus1 FRAP recovery curves normalized to the maximal recovery value. The mean recovery half-time and standard deviation is indicated. N=2 experiments, with n>9 cells each (n>46 cells in total). Shaded area shows the standard error. **G**. Boxplot of Myo52 fluorescence intensity at the fusion focus at fusion time. Bars are 5µm. Black or white p-values aligned with bars are relative to WT; grey ones to Fus1^Δ492-791^, p-values between bars compare the two conditions.

Strikingly, IDR deletions impaired the ability of cells to fuse, in a manner dependent on deletion length: *fus1*^*Δ501-749*^ cells fused better than *fus1*^*Δ501-791*^, which themselves fused better than *fus1*^*Δ492-791*^. The fusion-defective alleles also induced lysis in a large fraction of mating pairs, suggesting inaccurate location of cell wall digestion (Figure 4A,C). We note that the mutant with the widest distribution was also the most lysis-prone, consistent with the idea that cell lysis results from the reduced spatial precision of cell wall digestion (Figure 4B-C). These data demonstrate a clear functional role in cell fusion for Fus1 IDR.

### Fus1 IDR can be functionally replaced by heterologous self-assembling domains

We hypothesised that the Fus1 IDR may be functionally swapped with heterologous domains known to self-assemble, either by LLPS mechanisms, or by light induced oligomerization (Figure 4A, last four columns). The low-complexity domain of the mammalian Fused-in-Sarcoma protein (FUS^LC^ below) forms liquid condensates *in vivo* and *in vitro*, which age into solid fibrillar hydrogels [22-26]. FUS self-interaction is regulated by phosphorylation, and a phosphomimetic version of FUS^LC^, FUS^12E^ reduces aggregation and forms more liquid structures [27]. CRY2^PHR^ is the light-sensitive domain from *Arabidopsis* CRYPTOCHROME 2 protein, widely used in optogenetics studies [28]. Upon blue light exposure, CRY2^PHR^ oligomerizes [29], a property exacerbated in the CRY2^olig^ mutant [30]. By replacing the endogenous Fus1 IDR in the fully fusion-deficient mutant Fus1^Δ492-791^ by either of these four domains, FUS^LC^, FUS^12E^, CRY2^PHR^ and CRY2^olig^, our aim was to cover a broad range in types and strengths of self-interaction.

Strikingly, replacement of Fus1 IDR by FUS^LC^ or FUS^12E^ produced a fully functional formin that formed a concentrated focus at the fusion site, recruited Myo52 and supported cell-cell fusion to WT levels (Figure 4A-C). Thus, the condensation properties of Fus1 IDR can be functionally replaced by a heterologous self-assembling domain, which confirms the function of Fus1 IDR in self-assembly. It also demonstrates that this region fulfils no other essential function.

Replacement of Fus1 IDR by CRY2^PHR^ or CRY2^olig^ produced formin proteins that were fusion-incompetent in the dark, like the *fus1*^*Δ492-791*^ mutant, but partly supported cell fusion in the light (Figure 4D). For unknown reason, CRY2^PHR^ or CRY2^olig^ suppressed the lysis phenotype of *fus1*^*Δ492-791*^ mutant, even in the dark (Figure 4D). In the light, these two constructs formed foci close to the fusion site, but several observations indicate that these foci are not as functional as the FUS or WT variants. First, in pairs that successfully fused, foci of CRY2 variants showed a higher local concentration at the time of fusion than FUS or WT variants (Figure 4E). Second, in cell pairs that failed to fuse, the foci were initially positioned at the cell-cell contact, but then detached and moved away in each partner cells (Figure 4A; Movie S2). Third, analysis of the half-times of FRAP experiments showed that the CRY2^PHR^ and CRY2^olig^ variants recovered more slowly than the FUS or WT variants. In fact, we observed a progressive increase in Fus1 FRAP half-times, in the order WT, FUS^12E^, FUS^LC^, CRY2^PHR^, CRY2^olig^, from about 9 s in the WT to about 26 s in the CRY2^olig^ variant (Figure 4F). This is in agreement with previous work showing that CRY2^olig^ oligomerizes more strongly than CRY2^PHR^ and that FUS^LC^ forms a more solid phase than FUS^12E^ [27, 30]. These results agree with the view that CRY2^PHR^ and CRY2^olig^-containing Fus1 form more solid, less functional foci.

Finally, we observed that the Myo52 maximal intensities at fusion time were inversely correlated with the chimeric Fus1 mobilities measured by FRAP half times (Figure 4G). Note that these values, measured in successfully fusing cells, overestimate the recruitment of Myo52, which was often absent in CRY2-containing foci in unsuccessful mating pairs (Figure 4A). A likely explanation for this anticorrelation is that excessive Fus1 aggregation increases molecular crowding and focus solidity, thus impeding the entry of other proteins, including Myo52 and the vesicles it carries, in the structure. Together, these data indicate that the condensation properties of Fus1 are carefully tuned to yield a functional, selectively permeable fusion focus.

The gradual loss of clustering and function upon progressive deletion of Fus1 IDR suggests it supports weak, multivalent interactions, similar to those exhibited by the FUS^LC^ domain. The strength of these interactions may drive a fluid condensation that is permeable to actin-assembly factors such as actin-profilin and permits access to localization factors and myosin-driven cargoes, forming a cluster of secretory vesicles at the correct location. By contrast, stronger aggregation that solidifies the structure, as in the CRY2 constructs, likely restricts permeability and access, leading to detachment and lack of function. It will be interesting to investigate how Fus1 condensation properties are regulated by potential post-translational modifications during sexual differentiation and influence pheromone-MAPK signalling and regulation of actin assembly at the fusion focus [8]. In fact, the condensation of Fus1 to a very high local concentration provides an explanation for why key mutations in the FH2 domain that abolish actin assembly *in vitro* (at lower concentrations) only partly compromise fusion focus assembly [21, 31].

The condensation properties of Fus1 formin are essential to yield a focus that concentrates secretory vesicles for local cell wall digestion coordinated with the partner cell. This is reminiscent of the role of the synapsin protein, which phase-separates to organise clusters of synaptic vesicles at neuronal synapses [32]. Synapsin also bundles and promotes the assembly of actin filaments [33]. A similar mechanism may take place in budding yeast, and likely other fungi, where the formin-binding polarisome factor Spa2 was recently shown to phase-separate [34], likely promoting formin-dependent actin assembly to concentrate secretory vesicles at growth sites. Thus, biomolecular condensation by scaffolds linking to linear actin filaments, or in fission yeast directly by the formin nucleating the structure, may be a general principle by which to organize the focusing of secretory vesicles.

## Acknowledgements

We thank Aleksander Vjestica, Boris Sieber, Sjoerd Seekles, Sajjita Saha and Alejandro Melero-Carrillo for careful reading of the manuscript. This work was funded by grants from the Swiss National Science Foundation (310030B_176396 and 310030_191990) and the European Research Council (CoG CellFusion).

## Author contributions

IBC and SGM conceived the project. SGM performed the experiments in Figure 1D. OM performed the experiments in Figure 3. IBC performed all other experiments with technical assistance from LM. SGM acquired funding and coordinated the project. IBC and SGM wrote the first draft of the manuscript, which was revised by all authors.

## Materials and methods

### Strains construction

Strains were constructed using standard genetic manipulation of *S. pombe* either by tetrad dissection or transformation and can be found in Table S1. Oligonucleotides and plasmids used can be found in Tables S2 and S3, with details on how the plasmids were constructed.

*myo52-tdTomato:natMX, fus1-sfGFP:kanMX* and *tea1-mCherry:kanMX* tags were constructed by PCR-based gene targeting of a fragment from a template pFA6a plasmid containing the appropriate tag and resistance cassette, amplified with primers carrying 5’ extensions corresponding to the last 78 coding nucleotides of the ORF and the first 78 nucleotides of the 3’UTR, which was transformed and integrated in the genome by homologous recombination, as previously described [35]. Similarly, *acp2Δ::bleMX* was constructed by PCR-based gene targeting of a fragment from a template pFA6a plasmid containing the appropriate resistance cassette, amplified with primers carrying 5’ extensions corresponding or the last 78 nucleotides of the 5’UTR and the first 78 nucleotides of the 3’UTR, which was transformed and integrated in the genome by homologous recombination.

Construction of the strains expressing formin constructs from the *fus1* promotor at the *ura4* locus as a multicopy integration (Figure 1: *ura4-294:p*^*fus1*^*-fus1N*^*1-792*^*-fus1C*^*793-1372*^*-sfGFP:ura4+, ura4-294:p*^*fus1*^*-cdc12N*^*1-887*^*-fus1C*^*793-1372*^*-sfGFP:ura4+, ura4-294:p*^*fus1*^*-for3N*^*1-714*^*-fus1C*^*793-1372*^*-sfGFP:ura4+, ura4-294:p*^*fus1*^*:fus1N*^*1-792*^*-sfGFP:ura4+*) was done by homologous recombination of a transformed ura4^EndORF^-ura4^3’UTR^-p^fus1^-ForminConstruct-sfGFP-ura4^StartORF^-ura4^5’UTR^ fragment, obtained from StuI digestion of a pRIP based plasmid (pSM1659, pSM1663, pSM1662 and pSM1650, respectively). Such recombination recreates a new integration site, which has been shown to be unstable and to lead to multiple insertion [36], which is why we switched to single integration vectors for the rest of the study.

Construction of the strains expressing *fus1* constructs under *nmt1* promotor at the *ura4* locus as a single integration (Figure 2: *ura4+:p*^*nmt1*^*:fus1N*^*1-792*^*-sfGFP:termnmt, ura4+:p*^*nmt1*^*:fus1N*^*1-730*^*-sfGFP:term*^*nmt*^ *ura4+:p*^*nmt1*^*:fus1N*^*1-500*^*-sfGFP:term*^*nmt*^, *ura4+:p*^*nmt1*^*:fus1N*^*93-792*^*-sfGFP:term*^*nmt*^, *ura4+:p*^*nmt1*^*:fus1N*^*140-792*^*-sfGFP:term*^*nmt*^, *ura4+:p*^*nmt1*^*:fus1N*^*191-792*^*-sfGFP:term*^*nmt*^, *ura4+:p*^*nmt1*^*:fus1N*^*431-755*^*-sfGFP:term*^*nmt*^, *ura4+:p*^*nmt1*^*:fus1-sfGFP:term*^*nmt*^) was done by homologous recombination of a transformed ura4^5’UTR^-ura4^ORF^-ura4^3’UTR^-p^nmt1^-Fus1Construct-sfGFP-ura4^3’’^ fragment, obtained from PmeI digestion of a pUra4^PmeI^ based plasmid (pSM2600, pSM2601, pSM2644, pSM2630, pSM2825, pSM2703, pSM2645 and pSM2602, respectively). This leads to a stable single integration at the *ura4* locus [36].

Construction of the strains expressing tagged formin constructs from the endogenous locus (Figure 4: *fus1*^*Δ501-749*^*-sfGFP:kanMX, fus1*^*Δ501-791*^*-sfGFP:kanMX, fus1*^*Δ492-791*^*-sfGFP:kanMX, fus1*^*1-491*^*-FUS*^*12E*^*-fus1*^*792-1372*^*-sfGFP:kanMX, fus1*^*1-491*^*-FUS-fus1*^*792-1372*^*-sfGFP:kanMX, CRY2*^*PHR*^*-fus1*^*1-491*^*-fus1*^*792-1372*^*-sfGFP:kanMX, CRY2*^*olig*^*-fus1*^*1-491*^*-fus1*^*792-1372*^*-sfGFP:kanMX*) were done by homologous recombination of a transformed fus1^5’UTR^-ForminConstruct-sfGFP-kanMX-fus1^3’UTR^ fragment, obtained from a gel purified, SalI and SacII digested pFA6a based plasmid (pSM2507, pSM2697, pSM2625, pSM2941, pSM2940, pSM2937 and pSM2938, respectively). FUS fragments are the first 163 amino acids of the human protein. CRY2 fragments are the codon optimized PHR domains from *Arabidopsis thaliana. fus1Δ::LEU2+* [12] and *his5+:p*^*act1*^*:CRIB-3mCherry:bsdMX* [36] trace back to the aforementioned papers or are kind gifts from the afore mentioned labs.

### Growth Conditions

For mating experiments, homothallic (*h90*) strains able to switch mating types were used, where cells were grown in liquid or agar Minimum Sporulation Media (MSL), with or without nitrogen (+/-N) [37, 38]. For interphase experiments, cells were grown in liquid or agar Edinburgh minimal medium (EMM) supplemented with amino acids as required.

Live imaging of *S. pombe* mating cells was adapted from [38]. Briefly, cells were first pre-cultured overnight in MSL+N at 25°C, then diluted to OD600 = 0.05 into MSL+N at 25°C for 20 hours. Exponentially growing cells were then pelleted, washed in MSL-N by 3 rounds of centrifugation, and resuspended in MSL-N to an OD600 of 1.5. Cells were then grown 3 hours at 30°C to allow mating in liquid, added on 2% agarose MSL-N pads, and sealed with VALAP. We allowed the pads to rest for 30 min at 30°C before overnight imaging, or for 21h at 25°C for fusion efficiencies snapshot imaging, respectively.

For Correlative Light Electron Microscopy (CLEM) imaging, as described in [7], cells were grown for mating as described above or at exponential phase as described below. In the case of mating, after washes to remove nitrogen, cells were added into MSL™N plates. We allowed cells to mate for 5 h. A few microliters of MSL™N were pipetted onto the cells to form a thick slurry. In the second case, cells were pelleted by centrifugation. Yeast paste was pipetted onto a 3-mm-wide, 0.1-mm-deep specimen carrier (Wohlwend type A) closed with a flat lid (Wohlwend type B) for high-pressure freezing with a HPM100 (Leica Microsystems; for mating samples) or a Leica EM ICE high-pressure freezer (for interphase cells). The carrier sandwich was disassembled in liquid nitrogen before freeze substitution. High-pressure frozen samples were processed by freeze substitution and embedding in Lowicryl HM20 using the Leica AFS 2 robot as described [39]. 300-nm sections were cut with a diamond knife using a Leica Ultracut E or Ultracut UC7 ultramicrotome, collected in H2O, and picked up on carbon-coated 200-mesh copper grids (AGS160; Agar Scientific). For Light Microscopy, the grid was inverted onto a 1× PBS drop on a microscope coverslip, which was mounted onto a microscope slide. For Figure 3A-B, it was imaged using the DeltaVision platform described below to select for pairs with a Myo52-tdTomato and Fus1-sfGFP signal, indicating presence of a fusion focus. For Figure 3C-D, fluorescent TetraSpeck beads (Invitrogen), 100 nm in diameter, were adsorbed onto the grid before light microscopy imaging, to be used as fiducials for correlation. The grid was then imaged using the Zeiss LSM980 setup described below, to capture both Fus1N-sfGFP and fiducial fluorescence signal. The grid was then recovered, rinsed in H2O, and dried before post-staining with Reynolds lead citrate for 10 min. 15-nm protein A-coupled gold beads were adsorbed to the top of the section as fiducials for tomography.

For interphase imaging, cells were grown to exponential phase at 30°C in EMM+ALU media, pelleted and imaged between slide and coverslip. All strains containing a repressible *nmt* promotor were grown at least 24h without thiamine before imaging to reach maximal expression levels. For 1,6-hexanediol treatments in figures 2C, the drug was added directly before imaging to the final resuspension, to a final concentration of 20% and cells were imaged right away. For the 37°C treatment in Figure 2C, cells were grown to exponential phase at 30°C in EMM+ALU media, then shifted to 37°C for 6h, transported to the microscope on a 40°C carrier, and imaged at 40°C.

### Microscopy

Images presented in Figures 1B, 4A and S1A were obtained using a DeltaVision platform (Applied Precision) composed of a customized inverted microscope (IX-71; Olympus), a UPlan Apochromat 100×/1.4 NA oil objective, a camera (CoolSNAP HQ2; Photometrics or 4.2Mpx PrimeBSI sCMOS camera; Photometrics), and a color combined unit illuminator (Insight SSI 7; Social Science Insights). Images were acquired using softWoRx v4.1.2 software (Applied Precision). Images were acquired every 5 minutes during 9 to 15 hours. To limit photobleaching, overnight videos were captured by optical axis integration (OAI) imaging of a 4.6-μm z-section, which is essentially a real-time z-sweep.

Images presented in Figures 1D, 2B and 2C were obtained using a spinning-disk microscope composed of an inverted microscope (DMI4000B; Leica) equipped with an HCX Plan Apochromat 100×/1.46 NA oil objective and an UltraVIEW system (PerkinElmer; including a real-time confocal scanning head [CSU22; Yokagawa Electric Corporation], solid-state laser lines, and an electron-multiplying charge coupled device camera [C9100; Hamamatsu Photonics]). Time-lapse images were acquired at 1s interval using the Volocity software (PerkinElmer).

Images used to obtain Figures 2D and 4F were obtained using a ZEISS LSM 980 scanning confocal microscope with 4 confocal Detectors (2x GaAsP, 2x PMT), an Airyscan2 detector optimized for a 60x/1.518 NA oil objective, and 6 Laser Lines (405nm, 445nm, 488nm, 514nm, 561nm, 640nm) on inverted Microscope Axio Observer 7. For Figure 2D we used images acquired using the Airyscan2 detector and processed with the Zen3.3 (blue edition) software for super resolution. For Figure 4F we switched to the confocal mode as the lower fluorescence intensity required. We acquired images every second, and we bleached the cells by 1 iteration of a 25% (2D) or 10% (4F) 488nm laser power pulse after 5 time points and kept recording the fluorescence recovery for 5 (2D) or 2 minutes (4F). Temperature was controlled by an incubation chamber around the microscope.

Images used to obtain Figures 3A, 3B, 3E and 3F were obtained following CLEM, as described in [7]. TEMs were acquired on a FEI Tecnai 12 at 120 kV using a bottom mount FEI Eagle camera (4kx4k). Low-magnification tomograms were acquired at 4.576-nm pixel size and high magnification at 1.205-nm pixel size. For tomographic reconstruction of regions of interest, tilt series were acquired over a tilt range as large as possible up to ±60° at 1° increments using the Serial EM software [40]. The IMOD software package with gold fiducial alignment [41, 42] was used for tomogram reconstruction.

### Quantification and Statistical analysis

Percentages of cell fusion and lysis as in Figures 1C, 4C and 4D were calculated as in [6]. Briefly, 24h post-starvation, fused cell pairs, lysed pairs and the total number of cell pairs were quantified using the ImageJ Plugin ObjectJ, and percentages were calculated using the following equations:

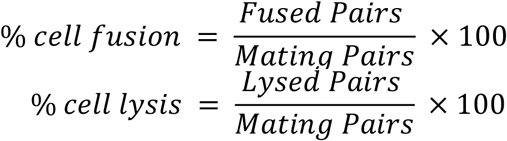

Fusion Focus intensities at fusion time as in Figure 1E, 4E and 4G were obtained from 5-minutes time lapse overnight movies using the maximum intensity of the Myo52-tdTomato dot to determine the moment of fusion (which correlates with the entry of GFP expressed under control of the P-cell-specific *p*^*map3*^ promoter into the *h-* partner [6]). On that time frame, a fluorescence profile across the fusion focus perpendicular to the long axis of the mating pair was recorded and either used directly as in Figure 1E or only the central point of the profiles were used to obtain boxplots as in Figure 4E and 4G. Profiles were background-subtracted and corrected for bleaching as follows: First, the cell fluorescence intensity was recorded over time in a square of about 7×7 pixels in 12 control (non-mating) cell. These fluorescence profiles were averaged, and the mean was fitted to a double exponential as it was describing our data better [43]:

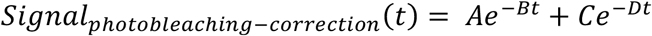

We then used this fit to correct the fluorescence profiles across the fusion focus for photobleaching. After subtracting background signal, the value at each timepoint was divided by the photo-bleaching correction signal:

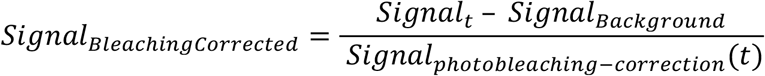

Corrected profiles were then either directly averaged and plotted (Figure 4E and 4G), or further normalized to the mean of the maximum (Figure 1E). Widths at half maximum (D50) as in Figures 1F and 4B were then calculated using these fluorescence profiles by recording (through linear interpolation in between points) the 2 distances that intersected with half maximal intensity for each individual profile, which were subtracted from one another. The result was then plotted as a boxplot.

The monopolar percentage as shown in Figure S1B was assessed from single fluorescence snapshot images of CRIB and classified as monopolar (decorating only one pole) or bipolar (decorating the two poles at similar intensities). The ratio of the first category divided by the sum of the two gave the monopolar percentage. Note that even WT bipolar cells can appear monopolar using this assay, as they can be captured at a time in CRIB oscillations where only one cell tip is decorated.

The density of vesicles in Figure 3C was obtained by manually counting vesicles within a half cylinder of 1µm diameter centred at the contact site.

The size of the ribosome free area as in Figure 3D was obtained by manually drawing the outline of the ribosome free area in each partner cell at the zone of cell-cell contact on one tomogram virtual slice and measuring its surface.

FRAP data analysis was performed by recording the fluorescence intensity of the bleached area using a manually fitted ROI, which was occasionally moved to track moving foci, which we could follow through the whole time-lapse as we only partially bleached the observed structures. Cells where the Fus1 foci could not be followed over the entire time course of the time lapse were excluded from the analysis. All the remaining traces were background substracted and bleach-corrected as above.

For Figure 2D, they were then scaled from minimum to pre-bleaching value as follows:

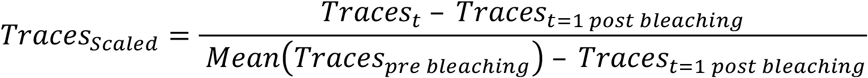

For figure 4F, they were scaled from minimum to maximum recovery value as follows :

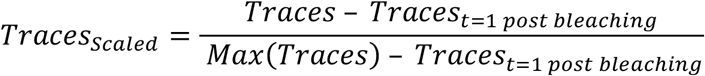

The resulting scaled traces were then averaged for each condition. These average traces were then used to fit the following conventional FRAP equation for each replicate and each condition:

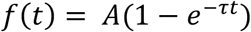

In the three replicates performed for Figure 2D, we obtained the following R^2^: 0.9844, 0.9917 and 0.9881 for Fus1N-Tips, 0.9534, 0.9683 and 0.96690 for Fus1N-Clusters. For the two replicates performed for Figure 4F, we obtained the following R^2^: 0.8836 and 0.9349 for WT, 0.9466 and 0.9349 for FUS^12E^, 0.9364 and 0.9869 for FUS, 0.9141 and 0.9677 for CRY2^PHR^ and 0.9135 and 0.9595 for CRY2^olig^. We used the fitted value of τ to calculate the half-time of recovery τ^½^ as follow :

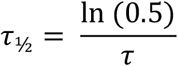

As we made respectively 2 or 3 replicates, we then obtained 2 (Figure 4F) or 3 (Figure 2D) values per condition, which were averaged and indicated directly on the figure along with their standard deviation. The graphs show the average from the first post-beaching point of all traces from all replicates for each condition along with their standard error.

Fiducial-based correlation was done using the Icy plug-in eC-CLEM [44], through 2D rigid transformation and manual matching of features. First correlation between light microscopy and low magnification electron microscopy images or tomograms was done using TetraSpeck beads, which are visible in both images. The resulting overlay images were then correlated with high magnification tomograms using 15-nm protein A–coupled gold beads as fiducials.

All plots, fittings, corrections and normalisations were made using MATLAB home-made scripts. For boxplots, the central line indicates the median, and the bottom and top edges of the box indicate the 25th and 75th percentiles, respectively. The whiskers extend to the most extreme data points not considered outliers. For bar plots, error bars represent the standard deviation. For the two FRAP plots, shaded areas represent the standard error. Statistical p-values were obtained using a two-sided student’s t-test, after normal distribution had been visually checked using a simple histogram. No further verification was made to ascertain that the data met assumptions of the statistical approach. All values below 0.05 are mentioned in the figures, including sample size.

---

**Figure S1 (related Figure 2).**
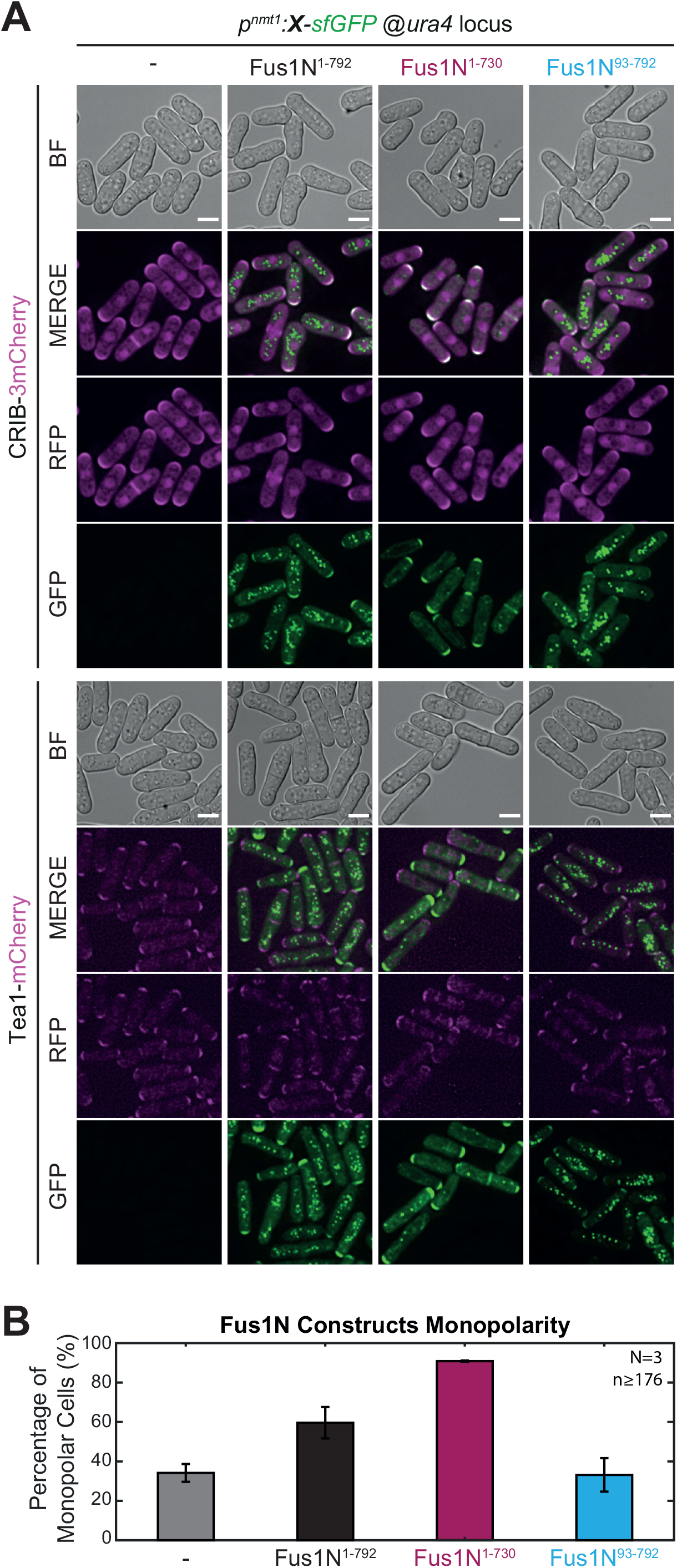
Fus1N tip localization induces monopolarity. **A**. DIC and fluorescence images of either (top) the polarity marker CRIB-3mCherry or (bottom) Tea1-mCherry in interphase WT or Fus1N^1-792^, Fus1N^1-730^ or Fus1N^93-792^-expressing cells. **B**. Monopolarity of the strains as in (A), assessed as the localization of CRIB on a single snapshot. Of note, a fraction of WT cells appear monopolar using this assay, because they are either before NETO or at a time point in CRIB oscillations [45] where only one tip is decorated. All p-values are relative to WT. Bars are 5µm.

## Movie legends

**Movie S1 (related to Figure 2). Interphase expression of Fus1 results in the formation of a fusion-focus like structure that polarize the cell at one cell tip and occasionally leads to lysis after division**

Fluorescence time lapse images of strains expressing full length Fus1-sfGFP from the *nmt1* promotor. The white arrowheads indicate cells that will lyse after division. Time is in hours:minutes. Bar is 5µm.

**Movie S2 (related to Figure 4). Aberrant behavior of fusion foci upon replacement of Fus1 IDR with CRY2**

DIC and GFP fluorescence time lapse images starting ∼4h post starvation of homothallic strains expressing either the WT fus1, or formin chimeras where the IDR has been replaced by either CRY2^PHR^ or CRY2^olig^, C-terminally tagged with sfGFP. The green and turquoise arrows mark cell pairs that have fused or not, respectively. Time is in hours:minutes. Bar is 5µm.

**Table S1:** Strains used in this study.

| GENOTYPE | FIGURES | STRAIN |
| --- | --- | --- |
| h90 myo52-tdTomato:natMX fus1-sfGFP:kanMX ura4- leu1-32 ade6-M216 | 1, 4 | YSM3312 |
| h90 myo52-tdTomato:natMX fus1Δ::LEU2+ ura4-294:p<sup>fus1</sup>-fus1N<sup>1-792</sup>-fus1C<sup>793-1372</sup>-sfGFP:ura4+ leu1-32 | 1 | YSM2504 |
| h90 myo52-tdTomato:natMX fus1Δ::LEU2+ ura4-294:p<sup>fus1</sup>-cdc12N<sup>1-887</sup>-fus1C<sup>793-1372</sup>-sfGFP:ura4+ leu1-32 | 1 | YSM2512 |
| h90 myo52-tdTomato:natMX fus1Δ::LEU2+ ura4-294:p<sup>fus1</sup>-for3N<sup>1-714</sup>-fus1C<sup>793-1372</sup>-sfGFP:ura4+ leu1-32 | 1 | YSM2510 |
| h90 myo52-tdTomato:natMX ura4-294:p<sup>fus1</sup>-fus1N-sfGFP:ura4+ fus1Δ::LEU2+ leu1-32 | 1 | YSM2486 |
| h90 myo52-tdTomato:natMX ura4-294:p<sup>fus1</sup>-fus1N-sfGFP:ura4+ leu1-32 | 1 | YSM2699 |
| h90 myo52-tdTomato:natMX ura4+:p<sup>nmt1</sup>-fus1N<sup>1-792</sup>-sfGFP:term<sup>nmt</sup> leu1-32 ade6-M210 | 2 | YSM4002 |
| h90 myo52-tdTomato:natMX ura4+:p<sup>nmt1</sup>-fus1N<sup>1-730</sup>-sfGFP:term<sup>nmt</sup> leu1-32 ade6-M210 | 2 | YSM4003 |
| h90 myo52-tdTomato:natMX ura4+:p<sup>nmt1</sup>-fus1N<sup>1-500</sup>-sfGFP:term<sup>nmt</sup> leu1-32 ade6-M210 | 2 | YSM4004 |
| h90 myo52-tdTomato:natMX ura4+:p<sup>nmt1</sup>-fus1N<sup>93-792</sup>-sfGFP:term<sup>nmt</sup> leu1-32 ade6-M210 | 2 | YSM4005 |
| h90 myo52-tdTomato:natMX ura4+:p<sup>nmt1</sup>-fus1N<sup>140-792</sup>-sfGFP:term<sup>nmt</sup> leu1-32 ade6-M210 | 2 | YSM4006 |
| h90 myo52-tdTomato:natMX ura4+:p<sup>nmt1</sup>-fus1N<sup>191-792</sup>-sfGFP:term<sup>nmt</sup> leu1-32 ade6-M210 | 2 | YSM4007 |
| h90 myo52-tdTomato:natMX ura4+:p<sup>nmt1</sup>-fus1N<sup>431-755</sup>-sfGFP:term<sup>nmt</sup> leu1-32 ade6-M210 | 2 | YSM4008 |
| h90 myo52-tdTomato:natMX ura4+:p<sup>nmt1</sup>-fus1-sfGFP:term<sup>nmt</sup> leu1-32 ade6-M210 | 2 | YSM4009 |
| h+ his5+:p<sup>act1</sup>:CRIB-3mCherry:bsdMX ura4-D18 | S1 | YSM4010 |
| h90 his5+:p<sup>act1</sup>:CRIB-3mCherry:bsdMX ura4+:p<sup>nmt1</sup>-fus1N<sup>1-792</sup>-sfGFP:term<sup>nmt</sup> leu1-32 | S1 | YSM4011 |
| h90 his5+:p<sup>act1</sup>:CRIB-3mCherry:bsdMX ura4+:p<sup>nmt1</sup>-fus1N<sup>1-730</sup>-sfGFP:term<sup>nmt</sup> ade6-M210 | S1 | YSM4012 |
| h90 his5+:p<sup>act1</sup>:CRIB-3mCherry:bsdMX ura4+:p<sup>nmt1</sup>-fus1N<sup>93-792</sup>-sfGFP:term<sup>nmt</sup> leu1-32 ade6-M210 | S1 | YSM4013 |
| h90 tea1-mCherry:kanMX ura4-D18 leu1-32 | S1 | YSM4014 |
| h90 tea1-mCherry:kanMX ura4+:p<sup>nmt1</sup>-fus1N<sup>1-792</sup>-sfGFP:term<sup>nmt</sup> leu1-32 | S1 | YSM4015 |
| h90 tea1-mCherry:kanMX ura4+:p<sup>nmt1</sup>-fus1N<sup>1-730</sup>-sfGFP:term<sup>nmt</sup> leu1-32 | S1 | YSM4016 |
| h90 tea1-mCherry:kanMX ura4+:p<sup>nmt1</sup>-fus1N<sup>93-792</sup>-sfGFP:term<sup>nmt</sup> leu1-32 ade6-M210 | S1 | YSM4017 |
| h90 myo52-tdTomato:natMX fus1-sfGFP:kanMX | 3 | YSM3888 |
| h90 myo52-tdTomato:natMX fus1-sfGFP:kanMX acp2Δ::bleMX ura4- leu1-32 ade6-M210 | 3 | YSM3314 |
| h90 myo52-tdTomato:natMX ura4+:p<sup>nmt1</sup>-fus1N<sup>93-792</sup>-sfGFP:term<sup>nmt</sup> fus1Δ::hphMX ade6-M210 leu1-32 | 3 | YSM4018 |
| h90 myo52-tdTomato:natMX fus1<sup>Δ501-749</sup>-sfGFP:kanMX ura4-294 leu1-32 ade6-M210 | 4 | YSM4019 |
| h90 myo52-tdTomato:natMX fus1<sup>Δ501-791</sup>-sfGFP:kanMX ura4-294 leu1-32 ade6-M210 | 4 | YSM4020 |
| h90 myo52-tdTomato:natMX fus1<sup>Δ492-791</sup>-sfGFP:kanMX ura4-294 leu1-32 ade6-M210 | 4 | YSM4021 |
| h90 myo52-tdTomato:natMX fus1<sup>1-491</sup>-FUS<sup>12E</sup>-fus1<sup>792-1372</sup>-sfGFP:kanMX ura4-294 leu1-32 ade6-M210 | 4 | YSM4022 |
| h90 myo52-tdTomato:natMX fus1<sup>1-491</sup>-FUS-fus1<sup>792-1372</sup>-sfGFP:kanMX ura4-294 leu1-32 ade6-M210 | 4 | YSM4023 |
| h90 myo52-tdTomato:natMX fus1<sup>1-491</sup>-CRY2PHR-fus1<sup>792-1372</sup>-sfGFP:kanMX ura4-294 leu1-32 ade6-M210 | 4 | YSM4024 |
| h90 myo52-tdTomato:natMX fus1<sup>1-491</sup>-CRY2olig-fus1<sup>792-1372</sup>-sfGFP:kanMX ura4-294 leu1-32 ade6-M210 | 4 | YSM4025 |

**Table S2:**
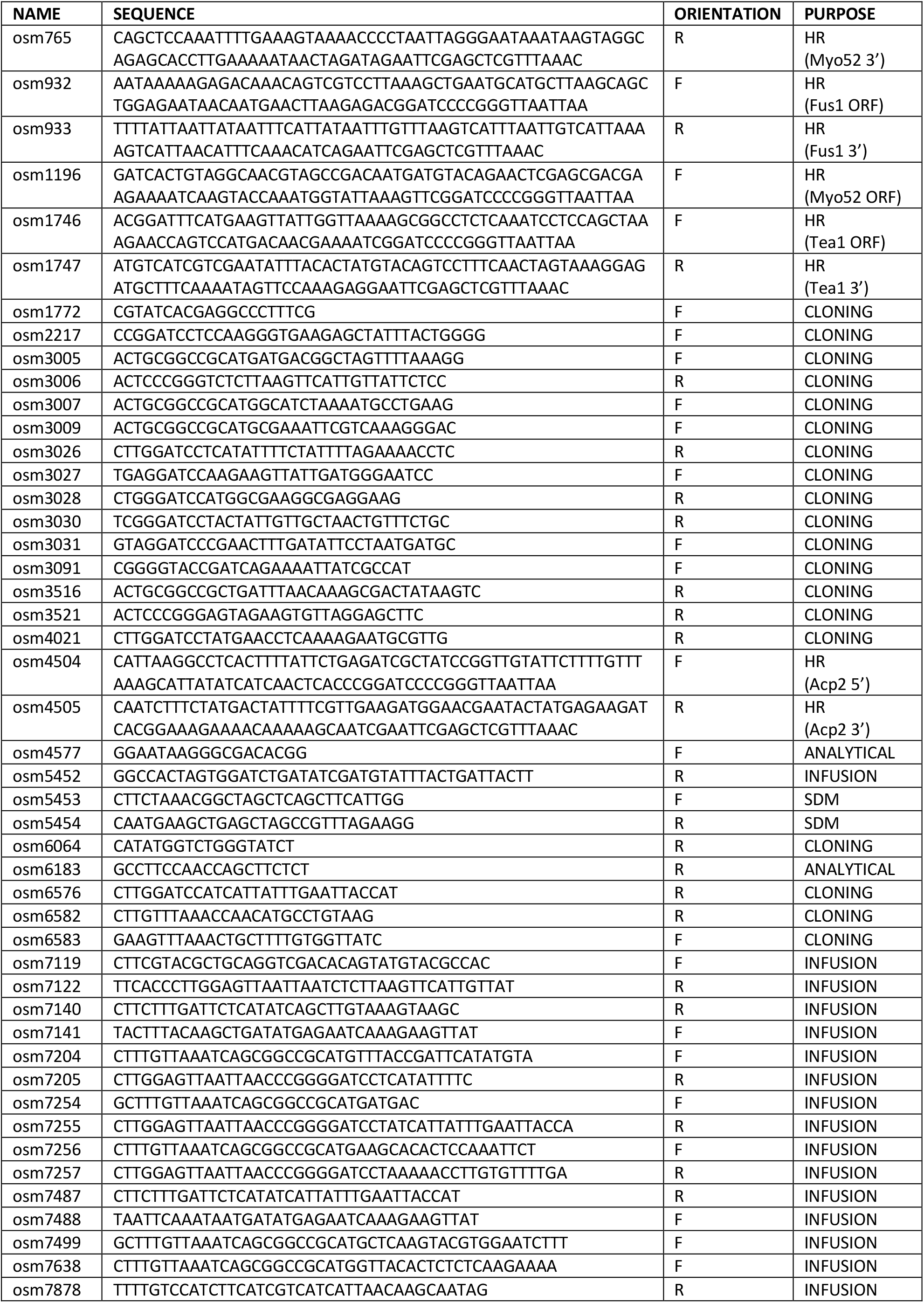
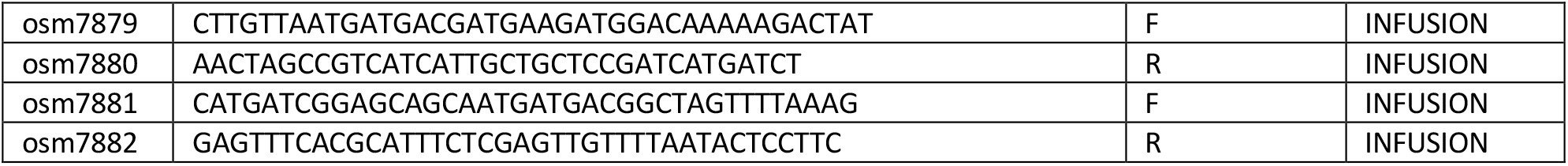
Primers used in this study. HR stands for homologous recombination in yeast [35] and SDM for site directed mutagenesis

**Table S3:** Plasmids used in this study. For each plasmid, the column “obtained from” indicates how it was constructed, from restriction enzyme-based cloning or infusion, with the primers and restriction enzymes used. “WT” indicates that genomic DNA from a wildtype strain was used as template for PCR amplification.

| NAME | DESCRIPTION | OBTAINED FROM | USAGE |
| --- | --- | --- | --- |
| pAV133 | pUra4<sup>AfeI</sup> | [36] | Single integration at *ura4* |
| pSM617 | pREP3x | Lab Stock | Pombe expression |
| pSM677 | pFA6a-mCherry-kanMX | Lab Stock | template for PCR-based HR |
| pSM685 | pFA6a-tdTomato-natMX | Lab Stock | template for PCR-based HR |
| pSM694 | pFA6a-bleMX | Lab Stock | template for PCR-based HR |
| pSM1538 | pFA6a-sfGFP-kanMX | Lab Stock | template for PCR-based HR |
| pSM1638 | pRIP-p<sup>fus1</sup>-sfGFP | Lab Stock | Multiple integration at *ura4* |
| pSM1650 | pRIP-p<sup>fus1</sup>-fus1N-sfGFP | Regular cloning :<br>pSM1638<sup>NotI/BamHI</sup>+(WT<sup>osm3005-osm3026</sup>)<sup>NotI/BamHI</sup> | Multiple integration at *ura4* |
| pSM1656 | pRIP-p<sup>fus1</sup>-fus1-sfGFP | Regular cloning :<br>pSM1638<sup>NotI/XmaI</sup>+(WT<sup>osm3005-osm3006</sup>)<sup>NotI/XmaI</sup> | Multiple integration at *ura4* |
| pSM1659 | pRIP-p<sup>fus1</sup>-fus1N-fus1C-sfGFP | 3-point ligation cloning :<br>pSM1638<sup>NotI/XmaI</sup>+(WT<sup>osm3005-osm3026</sup>)<sup>NotI/BamHI</sup><br>+(WT<sup>osm3027-osm3006</sup>)<sup>BamHI/XmaI</sup> | Multiple integration at *ura4* |
| pSM1662 | pRIP-p<sup>fus1</sup>-for3N-fus1C-sfGFP | 3-point ligation cloning :<br>pSM1638<sup>NotI/XmaI</sup>+(WT<sup>osm3007-osm3028</sup>)<sup>NotI/BamHI</sup><br>+(WT<sup>osm3027-osm3006</sup>)<sup>BamHI/XmaI</sup> | Multiple integration at *ura4* |
| pSM1663 | pRIP-p<sup>fus1</sup>-cdc12N-fus1C-sfGFP | 3-point ligation cloning :<br>pSM1638<sup>NotI/XmaI</sup>+(WT<sup>osm3009-osm3030</sup>)<sup>NotI/BamHI</sup><br>+(WT<sup>osm3027-osm3006</sup>)<sup>BamHI/XmaI</sup> | Multiple integration at *ura4* |
| pSM1823 | pRIP-p<sup>nmt41</sup>-sfGFP | Lab Stock | Multiple integration at *ura4* |
| pSM1826 | pRIP-p<sup>nmt41</sup>-fus1N-sfGFP | Subcloning :<br>pSM1650<sup>KpnI/NotI</sup>+pSM1823<sup>KpnI/NotI</sup> | Multiple integration at *ura4* |
| pSM2229 | pUra4<sup>AfeI</sup>-p<sup>nmt41</sup>-fus1-sfGFP | Lab Stock, Derived from pAV133 | Single integration at *ura4* |
| pSM2251 | pFA6a-fus1<sup>5'UTR</sup>-fus1_K879A-sfGFP-kanMX-fus1<sup>3'UTR</sup> | SDM :<br>pSM2827<sup>osm5453/5454</sup> | Single integration at endogenous *fus1* locus |
| pSM2390 | pRIP-p<sup>fus1</sup>-CRY2olig-For3N-fus1C-sfGFP | Lab Stock, Derived from pAV133 | Multiple integration at *ura4* |
| pSM2475 | pUra4<sup>AfeI</sup>-p<sup>fus1</sup>-CRY2PHR-fus1C-sfGFP | Lab Stock, Derived from pSM1662 | Single integration at *ura4* |
| pSM2478 | pUra4<sup>PmeI</sup>-p<sup>nmt41</sup>-fus1-sfGFP | 3-point ligation cloning :<br>pSM2229<sup>AatII/StuI</sup>+(pSM2229<sup>osm4577-osm6582</sup>)<sup>AatII/PmeI</sup>+(pSM2229<sup>osm6583-osm6183</sup>)<sup>PmeI/StuI</sup> | Single integration at *ura4* |
| pSM2507 | pFA6a-fus1<sup>5'UTR</sup>-fus1<sup>Δ501-749</sup>-sfGFP-kanMX-fus1<sup>3'UTR</sup> | 3-point ligation cloning :<br>pSM2251<sup>Sall/PacI</sup>+(pSM2251<sup>osm1772-osm6576</sup>)<sup>Sall/BamHI</sup>+(pSM2229<sup>osm3031-osm3521</sup>)<sup>BamHI/PacI</sup> | Single integration at endogenous *fus1* locus |
| pSM2600 | pUra4<sup>PmeI</sup>-p<sup>nmt1</sup>-fus1N-sfGFP | 3-point ligation cloning :<br>pSM2478<sup>KpnI/SacI</sup>+(pSM617<sup>osm3091-osm3516</sup>)<sup>KpnI/NotI</sup>+pSM1826<sup>NotI/SacI</sup> | Single integration at *ura4* |
| pSM2601 | pUra4<sup>PmeI</sup>-p<sup>nmt1</sup>-fus1N<sup>1-730</sup>-sfGFP | 3-point ligation cloning: | Single integration at *ura4* |
|  |  | pSM2600<sup>NotI/MscI</sup>+(pSM2600<sup>osm3005-osm4021</sup>)<sup>NotI/BamHI</sup>+( pSM2600<sup>osm2217-osm6064</sup>)<sup>BamHI/MscI</sup> |  |
| pSM2602 | pUra4<sup>PmeI</sup>-p<sup>nmt1</sup>-fus1-sfGFP | Subcloning :<br>pSM2600<sup>NotI/XmaI</sup>+pSM1656<sup>NotI/XmaI</sup> | Single integration at *ura4* |
| pSM2625 | pFA6a-fus1<sup>5'UTR</sup>-fus1<sup>Δ492-791</sup>-sfGFP-kanMX-fus1<sup>3'UTR</sup> | Infusion cloning :<br>pSM2507<sup>Sall/PacI</sup>+WT<sup>osm7119-osm7140</sup> +WT<sup>osm7141-osm7122</sup> | Single integration at endogenous *fus1* locus |
| pSM2630 | pUra4<sup>PmeI</sup>-p<sup>nmt1</sup>-fus1N<sup>93-792</sup>-sfGFP | Infusion cloning :<br>pSM2600<sup>NotI/XmaI</sup>+pSM2600<sup>osm7204-osm7205</sup> | Single integration at *ura4* |
| pSM2644 | pUra4<sup>PmeI</sup>-p<sup>nmt1</sup>-fus1N<sup>1-500</sup>-sfGFP | Infusion cloning :<br>pSM2600<sup>NotI/XmaI</sup>+pSM2600<sup>osm7254-osm7255</sup> | Single integration at *ura4* |
| pSM2645 | pUra4<sup>PmeI</sup>-p<sup>nmt1</sup>-fus1N<sup>431-755</sup>-sfGFP | Infusion cloning :<br>pSM2600<sup>NotI/XmaI</sup>+pSM2600<sup>osm7256-osm7257</sup> | Single integration at *ura4* |
| pSM2697 | pFA6a-fus1<sup>5'UTR</sup>-fus1<sup>Δ501-791</sup>-sfGFP-kanMX-fus1<sup>3'UTR</sup> | Infusion cloning :<br>pSM2507<sup>Sall/PacI</sup>+WT<sup>osm7119-osm7487</sup> +WT<sup>osm7488-osm7122</sup> | Single integration at endogenous *fus1* locus |
| pSM2703 | pUra4<sup>PmeI</sup>-p<sup>nmt1</sup>-fus1N<sup>191-792</sup>-sfGFP | Infusion cloning :<br>pSM2600<sup>NotI/XmaI</sup>+pSM2600<sup>osm7499-osm7205</sup> | Single integration at *ura4* |
| pSM2825 | pUra4<sup>PmeI</sup>-p<sup>nmt1</sup>-fus1N<sup>140-792</sup>-sfGFP | Infusion cloning :<br>pSM2600<sup>NotI/XmaI</sup>+pSM2600<sup>osm7638-osm7205</sup> | Single integration at *ura4* |
| pSM2827 | pFA6a-fus1<sup>5'UTR</sup>-fus1-sfGFP-kanMX-fus1<sup>3'UTR</sup> | Infusion cloning :<br>pSM1538<sup>Sall/EcoRV</sup>+IBC180<sup>osm7119-osm5452</sup> | Single integration at endogenous *fus1* locus |
| pSM2937 | pFA6a-fus1<sup>5'UTR</sup>-fus1<sup>1-491</sup>-CRY2PHR-fus1<sup>792-1372</sup>-sfGFP-kanMX-fus1<sup>3'UTR</sup> | Infusion cloning :<br>pSM2625<sup>Sall/XhoI</sup>+pSM2625<sup>osm7119-osm7878</sup> +pSM2475<sup>osm7879-osm7880</sup>+pSM2625<sup>osm7881-osm7882</sup> | Single integration at endogenous *fus1* locus |
| pSM2938 | pFA6a-fus1<sup>5'UTR</sup>-fus1<sup>1-491</sup>-CRY2olig-fus1<sup>792-1372</sup>-sfGFP-kanMX-fus1<sup>3'UTR</sup> | Infusion cloning :<br>pSM2625<sup>Sall/XhoI</sup>+pSM2625<sup>osm7119-osm7878</sup> +pSM2390<sup>osm7879-osm7880</sup>+pSM2625<sup>osm7881-osm7882</sup> | Single integration at endogenous *fus1* locus |
| pSM2940 | pFA6a-fus1<sup>5'UTR</sup>-fus1<sup>1-491</sup>-FUS-fus1<sup>792-1372</sup>-sfGFP-kanMX-fus1<sup>3'UTR</sup> | Infusion cloning :<br>pSM2625<sup>XhoI/SwaI</sup>+fus1<sup>XhoI site-491</sup>-FUS-fus1<sup>792-SwaI site</sup> ordered as a gBlock | Single integration at endogenous *fus1* locus |
| pSM2941 | pFA6a-fus1<sup>5'UTR</sup>-fus1<sup>1-491</sup>-FUS<sup>12E</sup>-fus1<sup>792-1372</sup>-sfGFP-kanMX-fus1<sup>3'UTR</sup> | Infusion cloning :<br>pSM2625<sup>XhoI/SwaI</sup>+fus1<sup>XhoI site-491</sup>-FUS<sup>12E</sup>-fus1<sup>792-SwaI site</sup> ordered as a gBlock | Single integration at endogenous *fus1* locus |

## Notes

### Competing Interest Statement

The authors have declared no competing interest.

